# SMaSH: A scalable, general marker gene identification framework for single-cell RNA sequencing and Spatial Transcriptomics

**DOI:** 10.1101/2021.04.08.438978

**Authors:** M. E. Nelson, S. G. Riva, A. Cvejic

## Abstract

Spatial transcriptomics is revolutionising the study of single-cell RNA and tissue-wide cell heterogeneity, but few robust methods connecting spatially resolved cells to so-called marker genes from single-cell RNA sequencing, which generate significant insight gleaned from spatial methods, exist. Here we present `SMaSH`, a general computational framework for extracting key marker genes from single-cell RNA sequencing data for spatial transcriptomics approaches. `SMaSH` extracts robust and biologically well-motivated marker genes, which characterise the given data-set better than existing and limited computational approaches for global marker gene calculation.

## Introduction

Single-cell RNA sequencing (scRNA-seq) [1; 2] is advancing our understanding of gene expression at the single-cell level in a variety of biological contexts. With scRNA-seq it is possible to study the multiplicity of both whole and partial transcripts in hundreds of thousands (and even millions) of individual cells, but there is no information on the location of different cell populations in tissue. Spatial transcriptomics addresses this issue by resolving the locations of the whole or part of the sequenced transcriptome. This additional spatial insight provides better context for studying the vast heterogeneity and interaction of different cellular states throughout different organs and tissues, making the integration of spatial and scRNA-seq analysis vital for gaining further insight to a variety of open problems in biomedical research. Several new rigorous analysis frameworks [3; 4] which integrate spatial transcriptomics and scRNA-seq data in a statistically robust manner have come online recently, and the merger of these technologies is gradually becoming standard practice in single-cell transcriptomics.

Broadly speaking spatial transcriptomics can be classified in ‘whole transcriptome’ and ‘specific transcript’ protocols. Whole transcriptome technologies, such as 10X Visium, allow the entire transcriptome to be resolved in tissue, but typically at the level of up to 10 cells per data point (a ‘spot’ on the Visium slice). In contrast, protocols such as seqFISH (sequential Fluorescence *In Situ* Hybridization) [5], ISS (*In Situ* Sequencing) [6], and MERFISH (Multiplexed Error-Robust FISH)^1^ [7] all aim to achieve this single-cell spatial resolution but for a limited number of genes.

Given the rich abundance and heterogeneity of gene expression across tissues, these approaches will come into their own only if the ‘right’ target genes can be accurately determined from the initial scRNA-seq data. Given that spatial transcriptomics resolves tissue at the single-cell level, and reveals multiple cell types across tissue compartments, it is important to select sets of genes which can be used to uncover both the global details of the tissue sample of interest and the local details of specific cell types and cell sub-types present therein. The challenge of selecting good markers is therefore complicated because it depends very much on the question the analyser cares about.

The scRNA-seq genes with expression profiles that are too noisy and/or highly-expressed across the bulk of the cell population will offer little to no insight from the tissue spatial analysis. Such expression patterns would be expected from e.g. housekeeping genes expressed throughout the tissue or genes with ribosomal or mitochondrial origins. At the same time, genes which have expression profiles which are too low will also be poorly resolved in the tissue due to the experimental limitations of existing technology. We will refer to genes which provide good global and local expression in spatially-resolved tissue sections, without being overly expressed throughout the sample and therefore simply ‘noise’ (or indeed too lowly expressed for detection), as *marker genes*. The exact list of marker genes for relevance to spatial transcriptomics depends on the problem at hand: different markers will be relevant if we wish to understand the spatial differences between different environments of the same cell type (e.g. tumour vs. healthy patient) or we wish to distinguish a broad class of cell types in the same tissue environment. The interesting marker genes for spatial analysis must therefore be inferred from a computational analysis of the corresponding scRNA-seq data which is general enough to calculate different markers for different questions which could be posed from the same available tissue data. At present, no such general automated approaches for selecting marker genes exist in the literature; only methods which select global marker genes based on standard gene expression patterns in scRNA-seq data are observed.

Current computational models [8; 9] for extracting marker genes from scRNA-seq data are limited in their scope and not well-suited to applications within spatial transcriptomics. These approaches identify marker genes based only on their expression profiles throughout the tissue of interest, leading to marker genes with large global expressions. Such highly-expressed genes are ineffective at distinguishing different cell types in the same tissue or different tissue environments for the same cell type because of their generic nature. We also found that these tools did not generalise well across different data-sets, producing marker genes which characterise some data-sets moderately well, but very poorly in other cases. Such markers are therefore not capturing the important information describing the ground-truth annotations from which they were originally determined for a large number of different biological scenarios. We also noted a lack of direct usability in current approaches with respect to popular computational pipelines, such as `ScanPy` [10].

To address these shortcomings, we propose the `SMaSH` (Scalable Marker (gene) Signal Hunter) framework (Figure 1), which identifies the key marker genes from scRNA-seq data for a variety of different problems to suit the interests of the analysis. `SMaSH` is motivated by the use case of selecting important genes for designing probes in upstream spatial transcriptomics experiments, such as *in situ* sequencing padlock probes. As such techniques are now in the transcriptomics mainstream there should be a robust, scalable, standardised approach to determining relevant markers. `SMaSH` has been designed for speed, so that it can suitably scale up from relatively small scRNA-seq data-sets of several thousand cells, to multi-million cell atlases such as [11]. Such a need for fast and efficient marker gene identification will be vital as we move into the ‘big data’ realm of computational single-cell biology. We also observe that `SMaSH` is able to identify robust marker genes for main cell types in a particular data-set, but also for the variety of cellular states that these main cell types can occupy (typically over 30). The few existing approaches to global marker selection are severely limited in performance at this task. `SMaSH` has been fully-integrated into the `ScanPy` Python framework [10] based on `AnnData` objects, which is one of the most popular platforms for scRNA-seq analysis to emerge in recent years. We believe this integration gives `SMaSH` a further edge as an efficient, highly-deployable, general, and user-friendly tool for robust marker gene extraction in an Python-based scRNA-seq analysis pipeline. To provide further enhancements to speed, `SMaSH` can be implemented in both computer-processing unit (CPU) and graphics processing unit (GPU) ‘modes’, where the latter is relevant for analysing the ever-growing data-sets under consideration in single-cell transcriptomics commonly spanning over 10^6^ cells.

**Figure 1.**
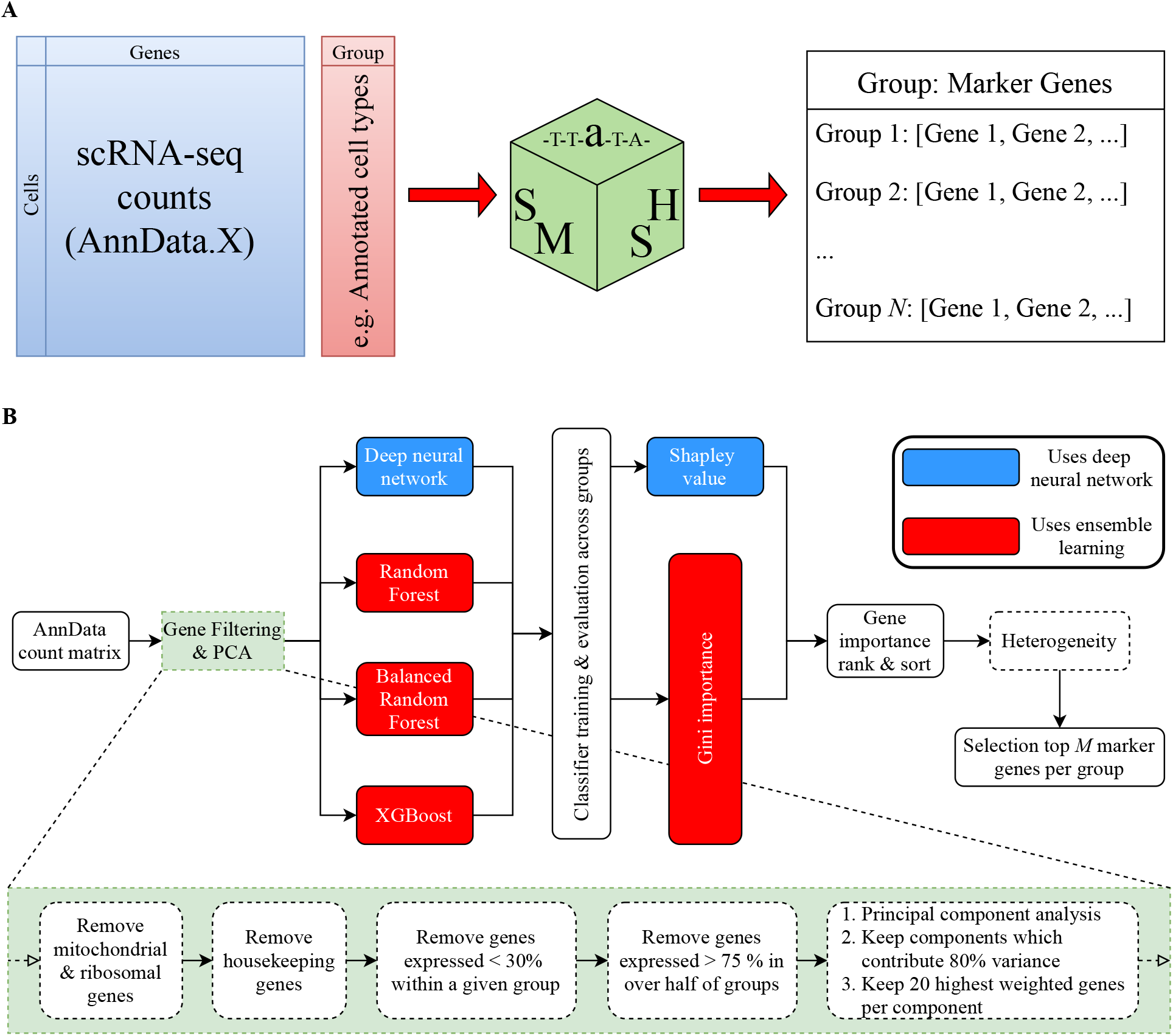
The SMaSH framework. **A)** `SMaSH` works directly from the counts matrix, produced a dictionary relating the user-defined classes of interest (e.g. cell type annotations) to top marker genes for each class (default top 5). **B)** `SMaSH` first filters out noisy and general genes, before keeping the those which contribute significantly to the final expression profile. These filtered genes are then ranked according to an ensemble learning model or a deep neural network, generating a final list of most important marker genes for each group or classification (e.g. different cell types) the user is interested in.

### SMaSH Framework

The `SMaSH` framework (Figure 1) is divided into four stages, beginning from the user-defined input `AnnData` [10] object which contains the raw scRNA-seq counts in a matrix of dimensionality determined by the number of barcoded cells and unique genes in the data-set. The user must also provide a vector of target outputs to map each barcoded cell onto, with values corresponding to classes depending on the problem in question. These can be, for example, a vector of annotations of each barcoded cell into a particular biological type, or the tissue or organ of origin of the cell, and so on. `SMaSH` then extracts markers by analysing the counts and targets in a supervised machine learning classification task, where the most important markers map to the most important features for classifying cells according to the user’s required targets. `SMaSH` is generic enough to calculate markers for any classification problem posed, provided the above conventions are adopted by the user. `SMaSH` implements its gene ranking and importance scoring using four different machine learning models: Random Forest (RF) [12], Balanced Random Forest (BRF) [13], `XGBoost` [14], and a feedforward deep feedforward neural network (DNN) [15; 16]. The first three models evaluate the importance score of each gene according to the *Gini importance* [17], whilst the neural network evaluates importance according to the *Shapley Value* [18]. Details of each step in the `SMaSH` framework are provided in the *Methods* section.

## Results

To evaluate the performance of `SMaSH`, we benchmarked it against two recent standalone computational al- gorithms, `scGeneFit` [8] and `RankCorr` [9], which calculate marker genes from scRNA-seq data using linear programming and gene-by-gene correlations respectively. Unlike `SMaSH` these algorithms determine relevant ‘global’ markers by considering the entire scRNA-seq counts matrix, and do not produce class-specific markers. We will demonstrate that, by using non-linear models to determine markers and class-specific marker calculations based on user-defined input annotations per cell, a more appropriate set of marker genes with improved classification of the bulk data can be achieved with `SMaSH`.

We compared `RankCorr`, `scGeneFit`, and each of the four models implemented within `SMaSH` across several publicly available data-sets: Zeisel [19], a data-set based on CITE-seq technology [20], a mouse brain single-nucleus RNA-sequenced data-set [4], a healthy foetal liver data-set [22], Paul15 stem cell data [21], and a large lung cancer data-set. We also considered an extension of the foetal liver data-set covering skin and kidney cells in addition to liver only when studying the performance of `SMaSH` on the problem of identifying organ-specific marker genes. Returning to the task of benchmarking `SMaSH`, the mouse brain data-set was split into two different sets of annotations, ‘broad’ and a higher-granularity where each broad cell type was further subdivided, in order to further study the effect of the cell annotation granularity on each of these data-sets. For the healthy foetal organ data-set, which spans the kidney, liver, femur, and yolk sack, we considered both the complete scRNA-seq data spanning all of those organs and the 40 different published annotations, and also separately the liver only where we applied our own set of cell annotations for that specific organ, corresponding to 18 different cell types. This was done to further study how the different marker gene frameworks responded to the same type of data but at different levels of complexity (18 distinct cell types vs. 40 in the full data-set). This 40 cell type data-set is an example of a particularly large data-set and we shall demonstrate the com- putational performance of `SMaSH` with respect to such a large ensemble of cells. These different data-sets use a variety of scRNA-seq technologies and conditions and were selected to give a cross-section of performance against both species and the cell size. The lung cancer data-set comprised non-small-cell lung cancer tissue, the 5 mm of tissue surrounding the tumour, and healthy lung tissue from donors. Annotations on this final data-set were performed using a combination of principal component analysis of the highly-variable genes for dimensionality reduction and manifold learning via UMAP [23] for visualisation purposes. These data-sets are summarised in Table 1. For the lung data annotations, as with mouse brain, there are two levels of complexity: first we defined seven ‘broad’ cell types corresponding to myeloid cells, B cells, T cells, dendritic cells, natural killer cells, mast cells, and epithelial cells. Each of these broad cell types, with the exception of epithelial cells, was then split into additional cell sub-types, resulting in 34 distinct classes in the final analysis. We will consider these two sets of annotations separately, again in order to study the performance of various models on the same data but with respect to different levels of complexity in the target assignment of the cells.

**Table 1.** Single-cell RNA-sequencing data-sets in this study. The different data-sets considered in the benchmarking of `SMaSH`.

| Data-set | Technology | Cells | Genes | # Cell types | Reference |
| --- | --- | --- | --- | --- | --- |
| Human lung cancer (broad) | 10X scRNA-seq | 54 574 | 18 612 | 7 | N.A. |
| Human lung cancer (cell sub-types) | 10X scRNA-seq | 54 574 | 18 612 | 34 | N.A. |
| Mouse brain (broad) | Single nucleus RNA-seq | 40 532 | 31 053 | 9 | [4] |
| Mouse brain (cell sub-types) | Single nucleus RNA-seq | 40 532 | 31 053 | 31 | [4] |
| Zeisel | 10X scRNA-seq | 3 005 | 4 000 | 7 | [19] |
| CITE-seq | CITE-seq | 8 617 | 500 | 13 | [20] |
| Paul15 | MARS-seq | 2 730 | 3 451 | 10 | [21] |
| Human foetal liver | 10X scRNA-seq | 65 712 | 19 572 | 18 | [22] |
| Human foetal organs | 10X scRNA-seq | 211 754 | 23 054 | 40 | [22] |

Each data-set has the form of an scRNA-seq matrix of UMI counts, with each row corresponding to a uniquely barcoded cell and each column a unique gene. There is also an associated vector of annotations for each cell corresponding to the experimentally determined cell type.

### Marker genes identifying broad cell types across different data-sets

In this first set of studies, we focused on the ‘broad’ cell types covering the broad human lung cancer, mouse brain, Zeisel, CITE-seq, Paul15, and human foetal liver; cell type multiplicities vary between 7 and 18. `scGeneFit`, `RankCorr`, and `SMaSH` separately calculated the most important 30 marker genes to classify cells according to their ground-truth annotations in each data-set. For each framework and data-set, the top 30 markers were then used as the only features in a *k*-nearest neighbours classifier for at mapping each cell back to its original annotation. The misclassification rates, *M*, and associated confusion matrices for recovering the original ground-truth annotations were evaluated on each data-set and model. The average *F*_1_ score was also calculated as the average harmonic mean of the precision and recall for each cell type classification, which is a more indicative performance metric for multi-classification problems than the more widely-known true-positive and false-negative rates. These performance metrics may be formally defined as:

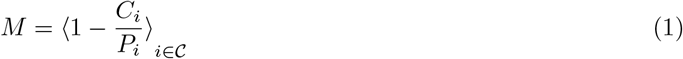

and

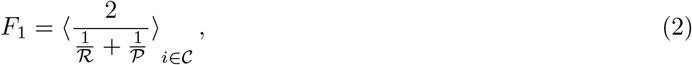

where *C*_*i*_ and *P*_*i*_ denote the number of correct predictions and total predictions of class *i* from the *k*-nearest neighbours classifier respectively, ℛ_*i*_ and 𝒫_*i*_ are the respective recall and precision of that classification, and the ⟨⟩_*i*∈𝒞_ denotes averaging over all classes *i* belonging to the set of annotations 𝒞 provided by the user. Lower misclassification rates (tending to 0) and higher average *F*_1_ scores (tending to 1) indicate better performance of a given model. The results are summarised in Table 2 for each framework and data-set.

**Table 2.** Marker gene misclassification rates in broad cell types. The average misclassification rates, *M*, in percent, and the weighted average *F*_1_ scores across all classes (cell types) for each data-set and framework, including the four different models implemented in `SMaSH`. All metrics are summarised as (*M, F*_1_) tuples. The top 2 performing models are indicated in bold red for each data-set. All `SMaSH` models outperform existing approaches across all data-sets.

| Data-set | scGeneFit | RankCorr | SMaSH (DNN) | SMaSH (RF) | SMaSH (BRF) | SMaSH (XGBoost) |
| --- | --- | --- | --- | --- | --- | --- |
| Human lung cancer (broad) | (25.5, 0.74) | (10.6, 0.89) | **(6.8, 0.93)** | (8.1, 0.92) | (9.6, 0.90) | **(7.4, 0.93)** |
| Mouse brain (broad) | (32.3, 0.67) | (8.6, 0.91) | **(0.4, 1.00)** | **(0.7, 0.90)** | (4.2, 0.95) | **(0.7, 0.99)** |
| Zeisel | (9.8, 0.90) | (7.3, 0.92) | (7.4, 0.92) | **(5.2, 0.95)** | (5.6, 0.94) | **(3.4, 0.97)** |
| CITE-seq | (17.8, 0.81) | (9.7, 0.89) | (7.5, 0.92) | **(7.1, 0.92)** | (13.4, 0.88) | **(7.2, 0.92)** |
| Paul15 | (27.8, 0.70) | (26.5, 0.73) | (25.6, 0.73) | **(19.4, 0.79)** | (34.4, 0.66) | **(14.5, 0.85)** |
| Human foetal liver | (57.8, 0.40) | (15.0, 0.85) | **(5.0, 0.95)** | **(4.9, 0.95)** | (8.5, 0.92) | (5.1, 0.95) |

We observe that the misclassification and general performance with `SMaSH` outperforms existing approaches across all data-sets, particularly for larger data-sets like the lung and human foetal liver, where `SMaSH` offers substantially lower misclassifications across all cell types. Thus, `SMaSH` scales very generally to marker gene identification problems in both simple data-sets like Zeisel and in larger data-sets, which are fast becoming the norm in single-cell biology. Confusion matrices of the true-positive (classification) rates for RankCorr, `scGeneFit` and the neural network and `XGBoost SMaSH` models evaluated on the ground-truth 7 broad cell types in the lung data are shown in Figure 2. We observe that, for both smaller and larger data-sets (e.g. Zeisel vs. broad lung) the ensemble learning and deep neural network models in `SMaSH` perform similarly. Performance of a given model varies with the data-set, and we would encourage all users to investigate several of the models available in `SMaSH` for their use case, but we note that `XGBoost` performs particularly well across all cases, and it the top two best performing models in 5/6 data-sets, and notably in the case of the mouse brain data achieve sub-percent misclassification rates where the best current approach of `RankCorr` achieve a 7.7 % average misclassification.

**Figure 2.**
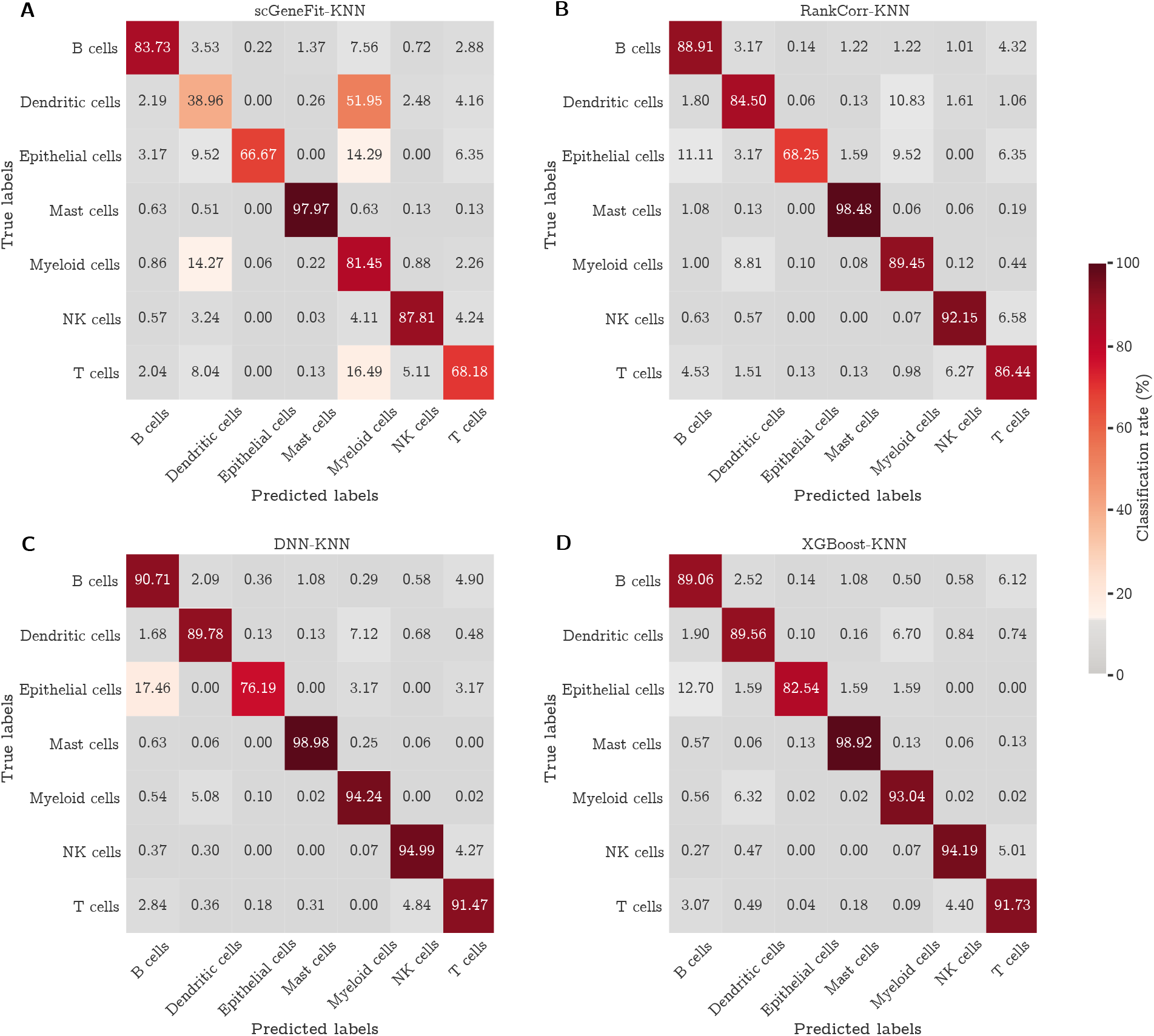
Lung broad cell type confusion matrices. Confusion matrices for the top 30 marker genes in the lung broad cell classification data-set, split by four different computational approaches to marker gene extraction: `scGeneFit` (A), `RankCorr` (B), `SMaSH` using the deep neural network (C), and `SMaSH` using `XGBoost` (D).

The `SMaSH` implementation provides the most important marker genes for each class, based on their rank in Gini importance or Shapley value. As a concrete example, in the case of the broad mouse brain data this would correspond to unique markers per each of the 9 cell types. These cell types biologically map to Astrocytes (Astro), Microglia (Micro), Endothelial cell (Endo), Excitatory neuron (Ext), Inhibitory neuron (Inh), Neuroblasts (Nb), Oligodendrocyte (Oligo), and Oligodendrocyte precursor (OPC), and a generic group of low quality cells (LowQ). These top three markers, ordered for each cell type based on their Shapley value computer by the deep neural network, are shown in Figure 3. In most cases, `SMaSH` is able to identify key genes which are uniquely (or nearly uniquely) expressed in one particular cell type of interest relative to all others. The colour scale, corresponding to the mean logarithm of gene expression, is normalised to between 0 and 1.0, where dark brown indicates very high levels of gene expression. Three dark brown populations can be uniquely generated for each cell type, indicating that highly and uniquely expressed genetic markers are present. Such markers would be useful for exclusively tagging particular cell types in the design of protocols for single-cell spatial resolution of mRNA e.g. the design of padlock probes for an *in situ* sequencing to use an earlier example. `SMaSH` is the first such technology we know of which automates marker selection per class in this fashion.

**Figure 3.**
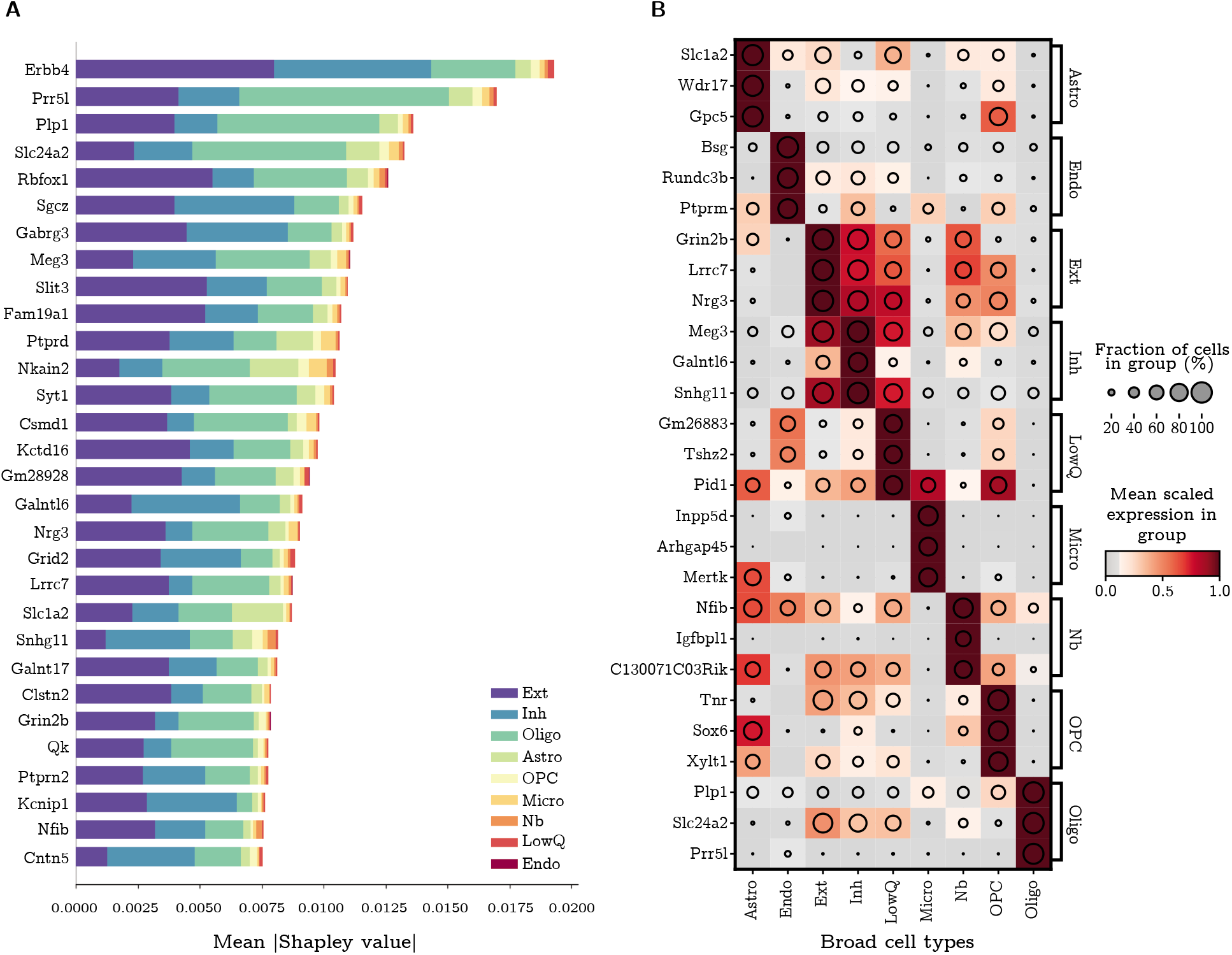
Marker genes for the broad mouse brain cell types. **A)** The mean (modulus) Shapley value for the top 30 ranked marker genes across all broad cell types of the mouse brain, before additional filtering and sorting, where the classification and marker extraction uses `SMaSH`’s deep neural network model. The Shapley value measures the average impact of that particular gene on the model, and the different colours indicate the different class contributions (where 0-8 label the 9 broad cell types of the mouse brain) which that particular gene explains. **B)** Following checks for heterogeneity of sorted markers, the final three markers for each class/broad cell type are shown, with the colour profile corresponding to the mean logarithm of the gene expression scaled between 0 and 1. The pattern of uniquely matching specific markers to specific cell types against all other cell types can be clearly seen as sets of three dark blocks (with maximal mean logarithm of gene expression) for each cell type. Shortened cell type names correspond to Astrocyte (Astro), Microglia (Micro), Endothelial cell (Endo), Excitatory neuron (Ext), Inhibitory neuron (Inh), Neuroblasts (Nb), Oligodendrocyte (Oligo), Oligodendrocyte precursors (OPC), and low miscellaneous low quality cells (LowQ).

### Marker genes identifying broad cell sub-types in lung cancer patients and mouse brain cells

One challenge in scRNA-seq gene identification is determining genes with the greatest statistical power for distinguishing increasingly complex and granular cell-type identifications. In the lung data-set each of the broad cell types can be further subclassified into several biologically distinct cell types. We repeated the misclassification calculation for 6 of the 7 broad lung cell types which can be further sub-divided, separately determining the top 30 markers for each of these 6 classification problems from the broad cell into its sub- types. We also evaluated this as a single classification problem, directly calculating the top 30 markers for classifying the entire lung data-set cells directly into their 34 lung cell sub-types. We evaluated `SMaSH` against existing approaches for identifying relevant markers, finding substantial reduction in misclassification rate compared to current methods. This was observed in both the ‘two-step’ approach of first classifying into the broad cell types, and then sub-classifying them, and the ‘one-step’ approach of directly classifying cells into the distinct 34 sub-types. We found that the misclassifcation rates for the ‘one-step’ problem were generally higher than the ‘two-step’ across all models. This is not unexpected given the added complexity of performing a 34-class problem directly and indicates that better marker gene extraction can be obtained by splitting the cell classification problem into two or more sub-problems. Moreover, we found that the largest gains in the ‘two-step’ problem are provided by either a more non-linear model, the deep neural network, or `XGBoost`. These comparisons are summarised in Table 3, where we also considered the same ‘one-step’ and ‘two-steps’ marker gene identification approach in the mouse brain data-set. Using `SMaSH` with a deep neural network and feature rankings based on the mean Shapley value of each gene, or `XGBoost` with a Gini importance ranking, greatly improved the ability to distinguish highly granular cell types. Both `scGeneFit` and `RankCorr` performed worse at this task. For the more complex ‘one-step’ classification `scGeneFit` and `RankCorr` do not perform well compared to any of the `SMaSH` models, and the neural network performs particularly well, benefiting from its ability to model and learn gene expression trends in a non-linear fashion.

**Table 3.** Marker gene misclassification rates in cell types in the lung and mouse brain. The average misclassification rates, *M*, in percent, and the weighted average *F*_1_ scores across all classes (cell types) for each human lung cancer cell sub-type and framework, including the four different models implemented in `SMaSH`. All metrics are summarised as (*M, F*_1_) tuples. The top 2 performing models are indicated in bold red for each data-set. All `SMaSH` models outperform existing approaches across all data-sets. HLC: Human lung cancer; MB: Mouse brain. Shortened mouse brain cell type names correspond to Excitatory neuron (Ext) and Inhibitory neuron (Inh), where well-defined sub-types could be extracted.

| Data-set | scGeneFit | RankCorr | SMaSH (DNN) | SMaSH (RF) | SMaSH (BRF) | SMaSH (XGBoost) |
| --- | --- | --- | --- | --- | --- | --- |
| HLC dendritic cell sub-types | (45.2, 0.53) | (6.9, 0.93) | **(5.6, 0.94)** | (5.7, 0.94) | (5.9, 0.94) | **(3.7, 0.96)** |
| HLC myeloid cell sub-types | (18.9, 0.80) | (13.8, 0.86) | **(11.9, 0.87)** | (12.4, 0.87) | (14.3, 0.85) | **(8.4, 0.91)** |
| HLC T cell sub-types | (25.6, 0.74) | (21.6, 0.78) | (20.0, 0.77) | **(16.2, 0.82)** | (20.8, 0.81) | **(19.7, 0.79)** |
| HLC B cell sub-types | (19.5, 0.79) | (7.8, 0.91) | (6.3, 0.93) | **(6.0, 0.93)** | (6.6, 0.92) | **(6.2, 0.93)** |
| HLC Mast cell sub-types | **(1.8, 0.98)** | (2.9, 0.97) | **(1.9, 0.98)** | (2.0, 0.98) | (2.0, 0.98) | (2.3, 0.97) |
| HLC natural killer cell sub-types | (14.3, 0.86) | (14.7, 0.85) | (12.6, 0.87) | **(11.4, 0.88)** | (18.1, 0.82) | **(11.8, 0.88)** |
| HLC all cell sub-types | (34.1, 0.65) | (34.0, 0.67) | **(16.9, 0.82)** | **(17.5, 0.82)** | (19.5, 0.80) | **(17.5, 0.82)** |
| MB Inh cell sub-types | (5.2, 0.95) | (4.3, 0.96) | **(1.7, 0.98)** | (5.6, 0.94) | **(1.7, 0.98)** | **(1.6, 0.98)** |
| MB Ext cell sub-types | (14.0, 0.86) | (14.2, 0.86) | **(4.2, 0.96)** | (4.8, 0.95) | **(4.6, 0.95)** | (5.1, 0.95) |
| MB all cell sub-types | (14.0, 0.86) | (21.1, 0.78) | **(3.9, 0.96)** | (5.2, 0.94) | (5.2, 0.95) | **(4.8, 0.95)** |

We also observe that `SMaSH` is still able to identify important marker genes which distinguish individual cell sub-types even when they belong to the same broad classification, as demonstrated for e.g. the sub-types of the mouse brain Inhibitory neuron broad types in Figure 4. For this Figure, the markers are calculated using the deep neural network in the case of `SMaSH` (C). The usual dark brown regions of high gene expression, now for a given cell sub-type, can be seen, and should be compared with the markers extracted from the using `scGeneFit` (A) and `RankCorr` (B). It can be seen that `SMaSH`, in addition to determining markers with lower misclassification rates, also produces markers which better represent each sub-type uniquely across the data. These results further support that, for a variety of data-sets and cell annotation complexities, `SMaSH` outperforms these current approaches in its ability to detect marker genes which almost uniquely capture the features of a particular cell type in both lower granularity (broad) and higher granularity (cell sub-type) annotation tasks.

**Figure 4.**
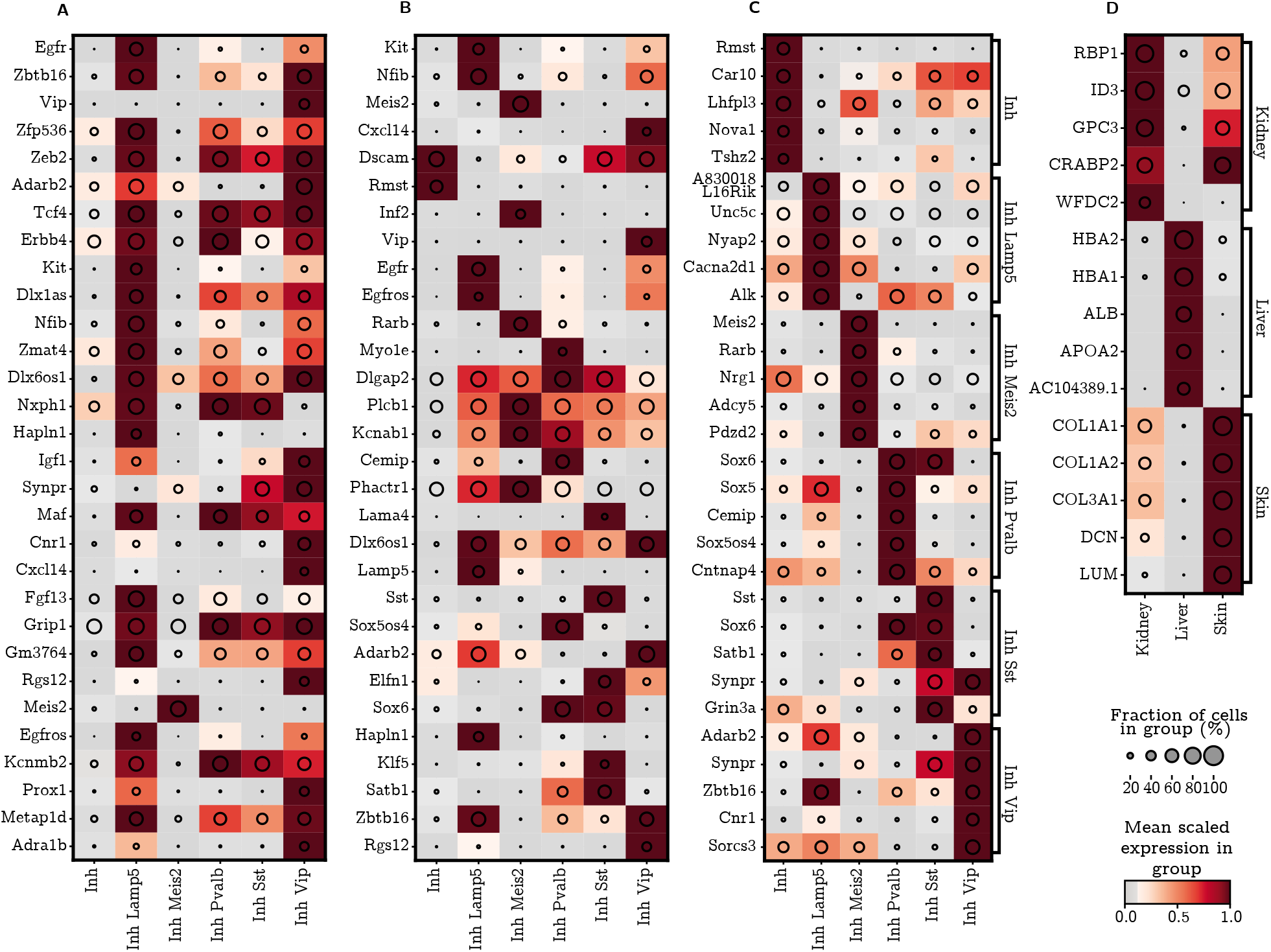
Marker genes for the mouse brain cell sub-types from the Inhibitory neuron broad types, and human foetal organ of origin classification. The mean logarithm of gene expression for mouse brain cell Inhibitory neuron cell sub-type markers. **A)** the markers for `scGeneFit`; **B)** the markers for `RankCorr`; **C)** markers from `SMaSH`’s default deep neural network model. Particularly in the case of `SMaSH` unique patterns can still be identified in this highly granular cell-type identification problem, whereas approaches such as `scGeneFit` are not able to identify many markers which uniquely resolve the sub-types present. **D)** `SMaSH` is able to select statistically significant markers for a highly imbalanced problem of distinguishing organs of origin in foetal scRNA-seq. Here the deep neural network, the usual default model, together with ranking based on Shapley values of genes, generates the final list of markers.

### Biological Interpretation of SMaSH

To investigate that `SMaSH` selects biologically versatile marker genes, we cross-checked the top markers per cell type and cell sub-type calculated for the mouse brain data across relevant literature. Table 4 summarises several example markers calculated with `SMaSH` for the broad cell types, their cell type and function, and existing references in literature, confirming that `SMaSH` correctly learns biologically robust and interesting lists of marker genes relevant to the underlying neurobiology. This list is far from exhaustive but the scope of the marker gene functions demonstrates that markers with a variety of biological functionality can be selected from a rich scRNA-seq data-set like the mouse brain.

**Table 4.**
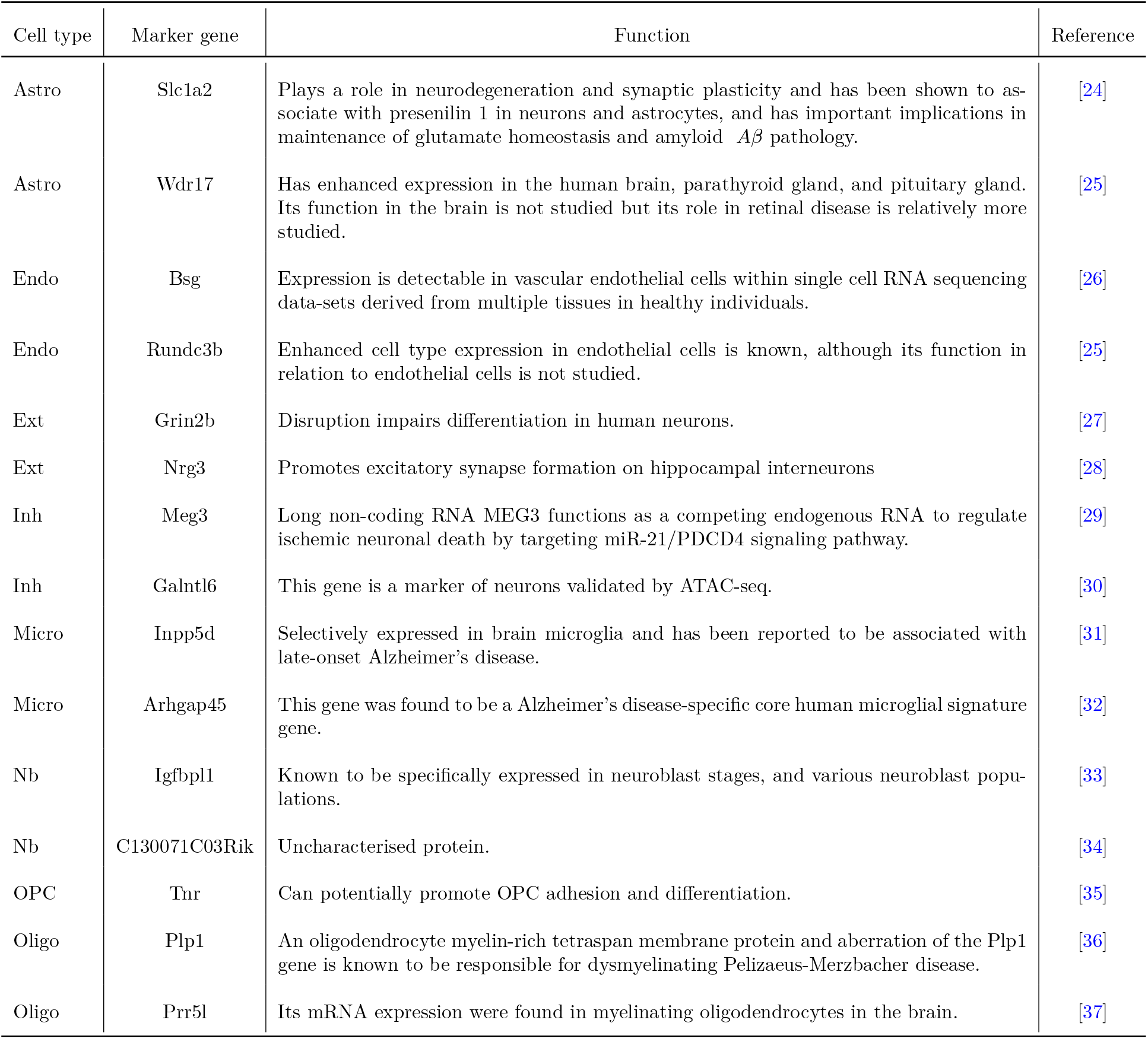
Marker gene functions for the broad mouse brain cells. Example markers genes across different broad cell types identified by `SMaSH`, together with known biological functions in the literature. Short- ened cell type names correspond to Astrocyte (Astro), Microglia (Micro), Endothelial cell (Endo), Excitatory neuron (Ext), Inhibitory neuron (Inh), Neuroblasts (Nb), Oligodendrocyte (Oligo), and Oligodendrocyte precursors (OPC).

### Marker genes differentiating organs of origin in early foetal development

In this section we demonstrate how `SMaSH` can be readily applied to very general marker gene identification problems in scRNA-seq. Thus far `SMaSH` has been implemented in problems for selecting marker genes to distinguish different cell types, which has obvious utility in spatial transcriptomics. However, this same procedure can be repeated in very general annotations and we illustrate this by taking a stratified sample of a publicly available foetal organ data-set [22] and calculating marker genes which best distinguish three different organs of origin: kidney, liver, and skin using those organs now as the relevant annotations for each cell. A similar problem would be e.g. distinguishing a tumour environment from healthy cells. Such identification problems are typically hindered by imbalanced data (e.g. many tumour samples but few healthy donors) and in the foetal organ case there are significantly more liver cells than kidney or skin cells in the scRNA-seq analysis [22]. In spite of these shortcomings, `SMaSH` is still able to identify statistically significant markers for specific organs, where the markers in question uniquely (or nearly uniquely) describe the particular organ of interest versus the other two in the classification problem (Figure 4**D)**). These markers were also confirmed to be highly relevant to the particular organ of interest following a cross-check of their function in relevant biological literature (Table 5). Given that an organ is a complex ensemble of many cell types, we may interpret an organ marker as a marker gene uniquely relevant to the function of dominant cell types for the organ of interest. For completeness we also benchmarked this problem against `scGeneFit` and `RankCorr` and find the lowest misclassification rate are achieved by `SMaSH`, with similar performance between the four models (Table 6), but with the best performance from the deep neural network and `XGBoost`, as observed in numerous other scenarios.

**Table 5.**
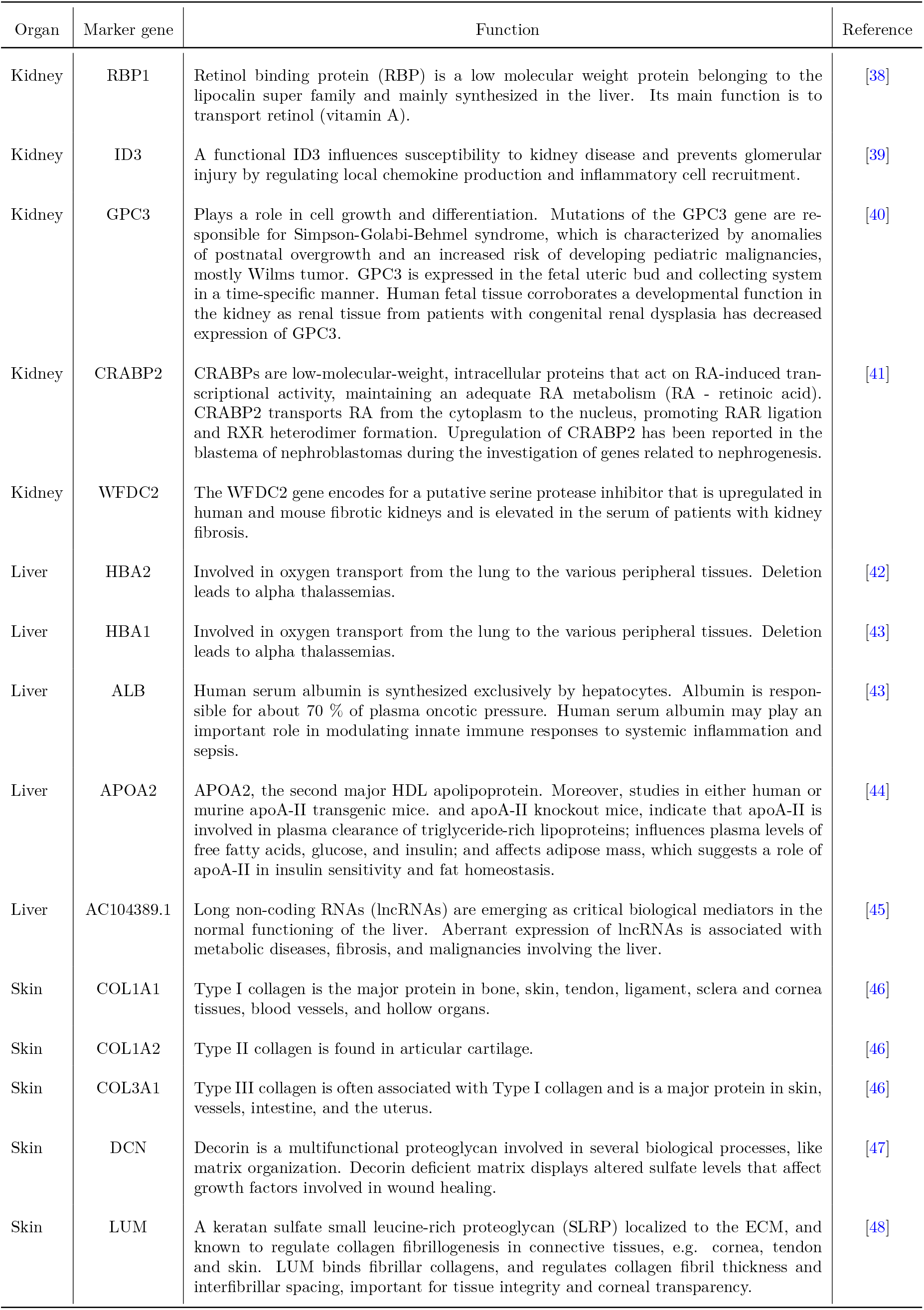
Marker gene functions for the classification of foetal organs. Example markers genes across different foetal organs (skin, liver, kidney) identified by `SMaSH`, together with known biological functions in the literature.

**Table 6.** Marker gene misclassification rates in organs of origin in early foetal development. The average misclassification rates, *M*, in percent, and the weighted average *F*_1_ scores across all classes (organs) in early foetal organ data, including the four different models implemented in `SMaSH`. All metrics are summarised as (*M, F*_1_) tuples. The top 2 performing models are indicated in bold red for each data-set. All `SMaSH` models outperform existing approaches across all data-sets. HFO: Human foetal organs.

| Data-set | scGeneFit | RankCorr | SMaSH (DNN) | SMaSH (RF) | SMaSH (BRF) | SMaSH (XGBoost) |
| --- | --- | --- | --- | --- | --- | --- |
| HFO skin vs. kidney vs. liver | (13.9, 0.85) | (5.2, 0.95) | **(1.1, 0.99)** | (1.4, 0.99) | (1.8, 0.98) | **(1.2, 0.99)** |

## Discussion

The `SMaSH` framework is a new methodology for determining marker genes from large scRNA-seq data-sets, for both and specific and general to a user-defined cell classifications (e.g a few broad cell types vs. many cell sub-types). This allows for more specific marker genes (e.g. markers for differentiating cell type A from cell type B) to be calculated in an automated and statistically robust fashion. To our knowledge, no such automated procedure exists for this purpose, so `SMaSH` was benchmarked against two recent approaches which determine ‘global’ marker genes across entire scRNA-seq counts matrices. We find that `SMaSH` produces markers which better classify data-sets of a variety of sizes and complexities, yielding markers which, when used to reconstruct the original annotations in each data-set, yield consistently lower misclassification rates. Such markers are therefore better able to uniquely classify the expression profiles of different cell types across these data-sets compared to the more global markers obtained in existing methods. This uniqueness applies to data-sets of varying granularity, as demonstrated by running `SMaSH` on separate human lung and mouse brain data-set in two modes: ‘broad’ cell classification of 7 different types for lung and 9 for mouse brain, and cell sub-types from each broad cluster leading to 34 distinct classifications of the lung cells and 31 distinct classifications for the mouse brain cells. Moreover, `SMaSH` is evaluated on data-sets with the variety of cells, ranging from 10^3^ to 10^5^, evaluating the markers in minutes. This makes `SMaSH` computationally tractable and scalable to high-throughput biological data-sets. `SMaSH` employs four different models which the user can specify, and it is recommended that the user study each of these models for the specific use case, but in general the performance of any model is substantially better than current approaches across most data-sets considered. In particularly, the performance of the deep neural network and `XGBoost` are consistently excel- lent in terms of yielding low marker gene misclassification rate in the data, high mean *F*_1_ score corresponding to high precision and recall in the marker extraction, and selecting final markers which allow for the visible distinction of cell types based on their mean gene expression profile. Therefore, combinations of these two models are recommended for the general use case. Markers are ranked based on explainability parameters which capture the information gain which each gene adds to the supervised model which aims to classify and reconstruct the user’s original annotation. In particular, we observe that ranking marker genes based on Shapley values is effective for revealing the most explainable features in the neural network model, and note that this measure explanability has yet to be explored in detail in applications of machine learning to problems in computational biological and transcriptomics.

`SMaSH` is available as a fully-integrated algorithm with `ScanPy`, making use of the `AnnData` object structure, common to many ‘big data’ analysis pipelines in single-cell computational biology. `SmaSH` is designed for robust marker gene identification across different cell types, and is specifically aimed for users wishing to identify marker genes relevant for wide varieties of different cell types which would be studied at the single cell resolution using specific spatial transcriptomics technologies. A notable example is for *in situ* sequencing, where 100-200 marker genes may be required for designing padlock probes which, when taken in combination, will attempt to spatially resolve the location of transcriptomes for the identification of both broad cell types and their cell sub-types in a variety of biological tissues and contexts. We summarise the `SMaSH` framework in a publicly-accessible webpage (see [pypi](https://pypi.org/project/smashpy/0.0.1/)), including self-contained notebooks where interested users can see example implementations for several data-sets mentioned in this paper (see GitLab repository). These materials demonstrate how a user may run `SMaSH` with any of its four models and obtain high-performance marker results consistent with what is documented in this paper. Based on this, we recommend `SMaSH` as a standard component to a downstream analysis pipeline of scRNA-seq data where key genes much be extracted, particularly with applications to spatial transcriptomics or related techniques in mind.

## Methods

### SMaSH Filtering Step 1: Gene Filter

The input cell-gene counts are first optionally batch-corrected using `Harmony` [49], and general genes connected to mitochondrial activity [50], ribosomal biogenesis [51], cell-surface protein regulation of the immune system, and biological housekeeping are removed. Genes which are lowly and highly expressed are further filtered out, so that only those which are expressed in greater than 30 % of the classes of interest and in less than 75 % of cells with more than 50 % of the classes of interest are retained. This final filter guards against additional batch-specific effects, such as a particular gene not being expressed uniformly across most various different independent biological samples comprising the data-set of interest.

### SMaSH Filtering Step 2: Inverse PCA

The filtered matrix of cells and genes is then dimensionally-reduced using principal component analysis (PCA) [52] applied to each gene as a unique feature. The PCA is then inverted and the top 20 genes in each principal component explaining up to 80 % of the overall variance in the data are retained. This additional feature guards against genes which would add very little extra information about the variance of expression profiles in the data and speeds up subsequent training of the model.

### Main SMaSH Model

The remaining genes are then ranked according to one of four machine learning classification models imple- mented in `SMaSH`: three ensemble learners (Random Forest (RF) [12], Balanced Random Forest (BRF) [13], and `XGBoost` [14] and a deep feedforward neural network (DNN) [15; 16]. Two different metrics are considered when ranking all genes in the problem according to how useful they are for classifying cells based on the initial target vector: the *Gini importance* [17] for ranking genes using the ensemble learners, and the *Shapley value* [18] for ranking genes with the neural network. The neural network model is used by default as it was found to provide very robust general performance with a lung data-set which motivated these initial studies. The neural network is a non-linear differentiable function and is therefore able to identify interesting non-linear patterns in the data related to gene expression, without resorting to simple ranking procedures which are linear, such as using correlation between genes.

The deep neural network is implemented with the `Keras` API [53], and its architecture determined by Bayesian hyperparameter optimisation with a Tree-structured Parzen estimator [54] as implemented in the `Hyperas` framework [55] where we started from a parsimonious set of different architectures. The network takes all filtered genes as unique input nodes, propagating weights through two hidden layers of 32 and 16 nodes respectively, each applying batch normalisation and sigmoid activation to the output. To aid regularisation, node drop-out is applied on a randomly selected 20 % and 10 % exiting the respective hidden layers. The user annotation classification is achieved by passing this output through a dense final layer of size equal to the number of unique annotations and a softmax function. Our decision to optimise over simple architectures is motivated by the observation that Shapley values are evaluated in exponential time, and therefore calculation of gene importance would slow down considerably using more complex architectures. The ensemble learners are implemented using the extensive `scikit-learn` library, and `imblearn` in the case of the BRF. Hyperparameter optimisation was performed using a greedy grid search across a large set of possible hyperparameters for each learner, selecting the parameters with the largest *F*_1_ score on the test-train split human lung cancer data-set, where a validation set from the initial testing set was selected to check for learning convergence and possible overtraining. The RF uses 100 estimators with no requirement on depth, and class weights are applied to the input data to reduce the impact of imbalanced samples. The BRF uses 200 estimators with a maximum depth of 50, and is also initialised with class weights and an ‘all’ sampling strategy. `XGBoost` uses 200 estimators with a maximum depth of 9, a softmax objective function (reducing to a logistic in the case of binary classification) with a 0.25 learning rate, and gradient boosting implemented through the `Dart` procedure.

### SMaSH Post-modelling: Ranking and Heterogeneity

The final marker genes are calculated by ranking and sorting the genes according to their total Gini importance or mean Shapley value, where the mean Shapley value is used by the default deep neural network. A set of relevant markers is produced for each class provided by the user from the initial vector of targets, where the top 5 markers per class are produced by default. A final heterogeneity check is conducted in the case that multiple samples are considered in the analysis, to make sure that the marker genes selected are also distributed uniformly in at least 70 % of the set of samples considered in the data. For this latter check the user must ensure that sample information is provided as an observation in the original `AnnData` object.

## Code Availability

The complete `SMaSH` implementation, including several full examples of how to use `SMaSH` and reproduce the results in the paper are available under the Cvejic group `GitLab`: https://gitlab.com/cvejic-group/smash.

## Data availability

We considered several publicly-available data-sets in this study: the Zeisel [19] data covering a small population of mouse brain cells, a data-set based on CITE-seq technology [20], a mouse brain single-nucleus RNA-sequenced data-set [4], a healthy foetal liver data-set [22], and Paul15 stem cell data [21]. We also considered an extension of the foetal liver data-set covering skin and kidney cells which is again available at [22]. These data-sets were modified from their originals to include the cell type annotations in a single `AnnData` object, for use directly with the tutorials to accompany the release of `SMaSH`. These modified data-sets are available on a self-contained google drive here.

## Lung Cancer Data-set

This data-set is based on lung tumour and non-tumourigenic tissues (background) collected from matched patients. Healthy tissues were also collected from deceased donors. The resulting cell suspensions were submitted for 3’ single cell RNA sequencing using Single Cell G Chip Kit, chemistry v3.1 (10X Genomics Pleasanton, CA, USA), following the manufacturer’s instructions. Libraries were sequenced on an Illumina NovaSeq S4, and mapped to the GRCh38 human reference genome using the Cell Ranger toolkit (version 3.0.0). Down- stream scRNA-seq analysis was then performed following the protocols implemented in the `ScanPy` workflow, with the final annotations determined manually and visualised using the Uniform Manifold Approximation and Projection (UMAP) space built from a neighbourhood graph applied to the top 15 principal components of the underlying gene space, and clustered using the Leiden algorithm based on graphical neighbourhoods.

## Acknowledgments

The authors wish to acknowledge Elisa Panda, Brynelle Myers, and Jiarui Xu for many helpful discussions on the biological interpretation of marker genes and its relevance to spatial transcriptomics, and Moritz Gerstung for advice on the preparation of the final manuscript and reviews of earlier drafts. The study was supported by European Research Council project 677501 – ZF_Blood (to AC) and a core support grant from the Wellcome Trust and MRC to the Wellcome Trust – Medical Research Council Cambridge Stem Cell Institute. MEN, SGR, and AC are further supported through the Open Targets project grant OTAR2060.

## Contributions

MEN conceived the study, developed the framework, wrote the manuscript, and supervised SGR; SGR developed and tested the framework, generated the results, and designed the final publicly-available tool; AC supervised SGR and wrote the manuscript.

## Footnotes

1 This list is far from exhaustive.

